# Multiomics profiling of APECED peripheral blood highlights compositional increases in alternatively activated B cell subsets

**DOI:** 10.64898/2026.07.24.740518

**Authors:** William Galbavy, Hyunjin Kim, Brian Klotz, Marine Malbec, Carley Tasker, Joseph C Devlin, Benjamin Daniel, Christina Adler, Wei Keat Lim, Asiel A. Benitez, Sokol Haxhinasto

**Affiliations:** Regeneron Pharmaceuticals, Tarrytown, New York, 10591. USA; Boehringer Ingelheim, Ridgefield, Connecticut, 06877, USA

## Abstract

APECED (Autoimmune PolyEndocrinopathy Candidiasis Ectodermal Dystrophy) is a rare syndrome of multi-organ autoimmunity driven by the presence of self-reactive T cells and autoantibodies caused by mutations in the gene Autoimmune regulator (*AIRE*). Compared to the well-defined role of AIRE in establishing and maintaining T cell central tolerance, less is known about how AIRE deficiency impacts B cell phenotypes that may contribute to a breakdown in peripheral B cell tolerance. Here we analyzed serum and peripheral blood cells from APECED patients and healthy donors using autoantibody profiling, proteomics, flow cytometry, scRNAseq based subset analysis, BCRseq, and autoantigen binding assays finding significant changes to the APECED B cell compartment. We show that while Naive and Transitional B cells are reduced, alternatively activated B cell subsets are expanded in APECED patients including IgM CD27+ and class switched Atypical B cells which exhibit BCR chain features prone to autoreactivity. Antibodies derived from either APECED or Healthy class switched atypical B cells bind autoantigens at significantly higher rates than control IgG B cells. Serum autoantibodies and proteomics highlight APECED common and patient variable changes. These results together show that APECED causes a compositional shift towards subsets that may promote broken B cell tolerance and autoimmunity.

## INTRODUCTION

Autoimmune PolyEndocrinopathy Candidiasis Ectodermal Dystrophy (APECED) is a rare autoimmune syndrome caused by mutations in the AutoImmune REgulator (AIRE) gene (1–6). AIRE deficiency disrupts T cell central tolerance, leading to the survival of autoreactive T cells and the production of autoantibodies which drive multi-organ autoimmunity (7, 8). While the role of AIRE in regulating T cell central tolerance via medullary thymic epithelial cells is well-established, the impact of AIRE deficiency on B cell peripheral tolerance remains underexplored (9–12).

B cells play significant roles in autoimmunity through their abilities to present antigens, traffic to organs, produce cytokines, and generate autoantibodies. In APECED disease B cell tolerance is broken, producing a wide variety of autoantibody species that are associated to clinical manifestations. In APECED patients, Type I IFN autoantibodies are present that cause worsened outcomes to viral infections including SARS-CoV2 (13, 14), influenza (15), and fulminant viral hepatitis (16). These autoantibodies can have neutralizing effects on most of the IFN alpha family cytokines (17) and downregulate the Type I IFN gene signature of immune cells in APECED (18). Autoantibodies to IL17 family cytokines associate to (19) and help drive (20, 21) chronic mucocutaneous candidiasis (CMC) along with the Th1 skewed immune response in APECED (22, 23). Other known biologically impactful autoantibodies in APECED include those against intrinsic factor that cause pernicious anemia, or those against tryptophan hydroxylase (TPH1) in the serotonin biosynthesis pathway are related to reduced serotonin levels and the presence of gastrointestinal symptoms (24). AIRE deficient mice produce autoantibodies (6, 7, 25) that vary by strain (26), and B cells are necessary to drive key autoimmune phenotypes including exocrine pancreatitis in the NOD background (10, 27). Around one hundred or more autoantibodies have been described in APECED patients (28, 29, 30). These autoantibody species contain both diagnostic potential for patients and serve as either known or potential drivers of autoimmunity (11). Despite this research activity surrounding the presence and function of autoantibodies in APECED, the mechanisms by which AIRE deficiency impacts B cell subsets and their propensity for autoreactivity remains poorly understood. Previous studies suggest AIRE deficiency may disrupt a peripheral tolerance checkpoint on B cells normally provided by a healthy and diverse T cell clonal repertoire, leading to the accumulation of an autoreactive-prone mature naïve B cell subset (9). Others have suggested that conditions in the lymph nodes specifically are dysregulated in APECED, leading to increased T follicular helper (Tfh) cells and smaller germinal centers that may be driving B cell autoimmunity through autoreactive Tfh mediated T cell help and faulty Tregs (31). Nevertheless, the specific B cell populations contributing to autoantibody production and their molecular characteristics in APECED patients have not been fully delineated. This study aims to address these gaps by investigating the composition, transcriptional signature, BCR phenotype, and autoreactive features of B cell subsets in a cohort APECED patients. Using single-cell multi-omics including scRNA/BCR sequencing and complementary autoantigen binding assays, we provide a high-resolution analysis of B cells in APECED.

We find that while Naïve and Transitional B cells are decreased, alternatively activated IgM CD27+ and atypical B cell subsets are enriched in the APECED B cell compartment and that antibodies derived from class switched atypical B cells as a cell type are more likely to bind autoantigens than antibodies of control IgG B cells.

Serum proteomic profiling contributes to this picture, showing that APECED patients have increases in CXCL9, reductions in IL22, and an overall pro-inflammatory signature including interferon gamma inducible proteins, acute phase proteins, and those associated to the innate immune system. Our findings reveal significant shifts in B cell composition, implicating decreases to Naïve populations and increases to IgM CD27+ and class switched atypical B cells with autoreactive prone BCR features as potential contributors towards the breakdown of peripheral B cell tolerance in APECED.

## RESULTS

### Autoantibody profiling identifies high levels of autoantibodies against diverse autoantigens in sera from from APECED patients

To establish and survey the extent of broken B cell tolerance through the production of autoantibodies in our patient cohort, we first tested APECED patient serum for the presence of anti-cytokine autoantibodies. We detected the presence of significantly higher IgG signals against IFN-w, IFN-A2, IL17F, IL17A, IL22, IL6, and IFN-b cytokines from APECED serums compared to Healthy serums (Figure 1A). Autoantibodies against the type I IFNs are highly prevalent (11, 12), used clinically to aid in diagnosing APECED disease (especially IFN-w), and were observed in every patient (10/10 patients). IL17F (9/10 patients), and IL22 (10/10 patients) autoantibodies are also observed at a high rate in APECED patients while IL17A is common but not as prevalent (19, 20, 21) and observed in fewer of our patient sera. IL6 and IFN-b autoantibodies are less common but have been detected in APECED (32, 18).

**Figure 1.**
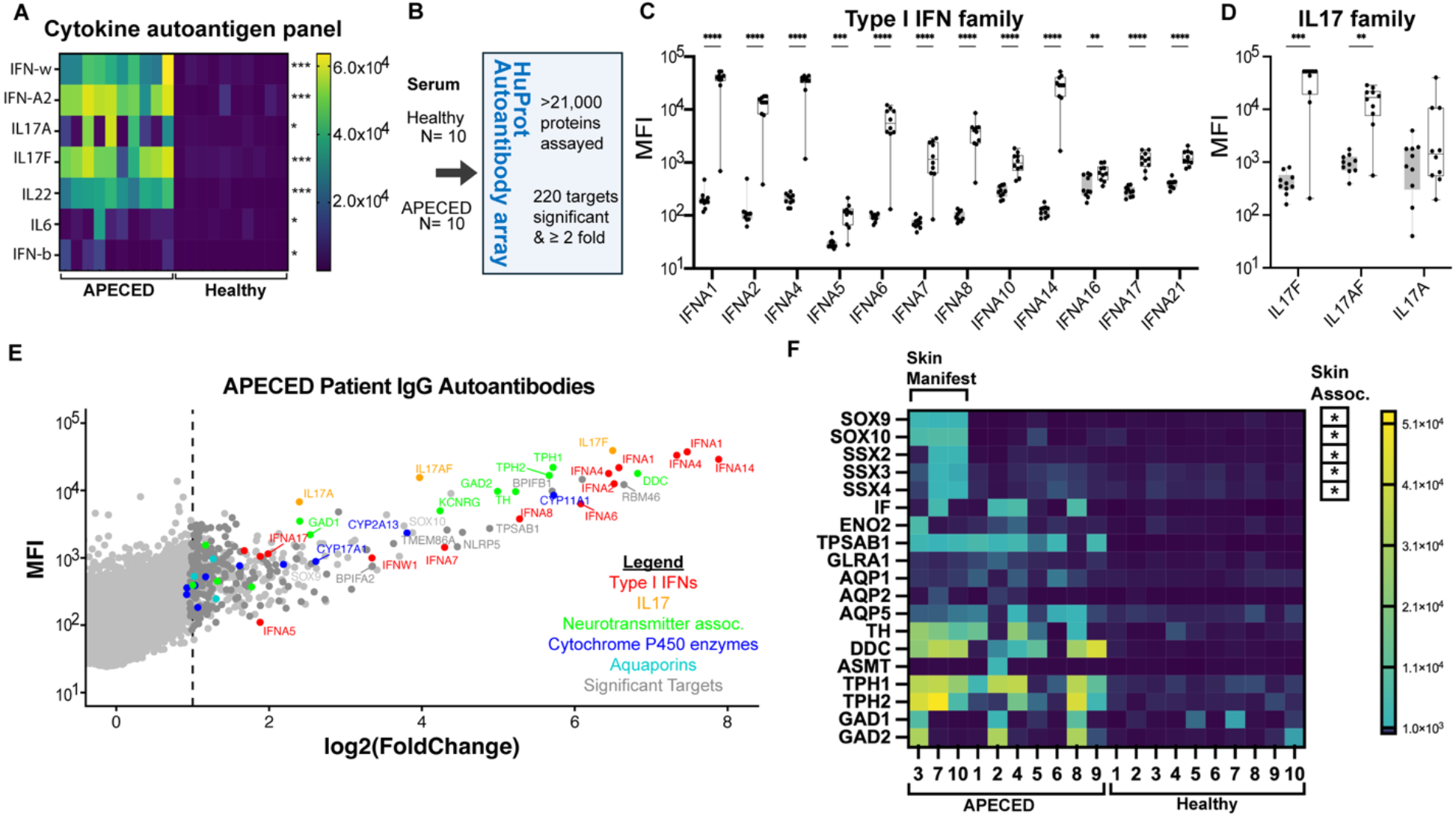
Autoantibody profiling identifies high levels of autoantibodies against diverse autoantigens in sera from APECED patients. (A) Anti-cytokine panel for APECED and Healthy donor IgG autoantibodies. Wilcoxon rank-sum tests * p < 0.05 *** p < 0.001. (B) HuProt array IgG autoantibody profiling from patient serum defined 220 targets with MFIs statistically significant between groups by Wilcoxon rank sum test and enriched by ≧ 2 fold in APECED over Healthy Control. (C-E) Detection of major anti-cytokine autoantibodies in APECED. MFI values per patient for (C) Anti Type I IFNs, (D) anti-IL17. (E) Plot of group enriched autoantibodies in APECED vs Healthy control samples. Average APECED autoantibody MFI signal plotted with average Log2 fold change over Healthy control. (F) Individual variability of APECED patients in select autoantibody MFIs. SOX9, SOX10, SSX2, SSX3, and SSX4 MFIs are increased in APECED patients with skin manifestations compared to APECED patients without skin manifestations (Wilcoxon rank sum tests * p < 0.05).

As previous reports have noted, APECED patients can have a diverse range of autoantibodies that have the potential to impact biological processes. In order to assay the wider extent of autoantibody production in APECED patient serum, we then used an autoantibody profiling array (HuProt Array, CDI) of >21,000 proteins to uncover ∼200 APECED group enriched autoantibody targets (Figure 1, B-F, see supplemental spreadsheet).

Consistent with prior reports (28, 29, 30), autoantibodies against Type I interferon family cytokines (IFNA1, IFNA2, IFNA4, IFNA5, IFNA6, IFNA7, IFNA8, IFNA10, IFNA14, IFNA16, IFNA17, IFNA21), IL17 family cytokines (IL17F, IL17AF, IL17A), Cytochrome P450 enzymes (CYP2A6, CYP11A1, CYP2A7, CYP2B6, CYP2D6, CYP4F11, CYP2A13, CYP1A1), and various neurotransmission associated enzymes (TPH1, TPH2, DDC, TH, GAD2, ASMT) were among the autoantibodies observed in our APECED patient cohort (Figure 1, B–D). Surprisingly, the IL22 protein included in the HuProt autoantibody array did not yield any IgG signal despite detecting anti-IL22 autoantibodies in every APECED patient’s serum across multiple orthogonal assays. It is unclear why, but it may be due to protein misfolding caused by the GST tag.

Interestingly, antibodies against two transcription factors essential for melanocyte development SOX9 and SOX10 were found significantly increased (SOX9 p = 0.0167, SOX10 p = 0.0167) in patients with skin manifestations (patients 3, 7, 10) including vitiligo (patients #7 and #10), a phenomenon documented previously in APECED patients (33). We additionally found antibodies targeting synovial sarcoma X chromosome breakpoint (SSX) proteins SSX2, SSX3, and SSX4 with unknown functionality enriched in APECED patients with skin manifestations (SSX2 p = 0.033, SSX3 p = 0.0167, SSX4 p = 0.033) including vitiligo (Figure 1D). SSX proteins such as SSX2 are found expressed in cancers such as synovial sarcoma, melanoma, and others but they are specific for testis in normal tissues (34). Anti-intrinsic factor antibodies were observed in four APECED patients (Figure 1D) and have been associated with gastritis and pernicious anemia (35). APECED patients 1, 2, and 4 were noted to have anemia, anti-IF antibodies were observed in two of three of these patients.

Autoantibodies targeting the aquaporin family including AQP5 were found in subsets of APECED patients. Other APECED enriched IgG autoantibody targets of interest were against mast cell alpha beta tryptase TPSAB1, the glycine receptor subunit GLRA1, and neuron specific enolase ENO2. These autoantibody species have been previously observed in Sjogrens (36), Rheumatoid Arthritis (37), motor disorders (38), and eye disease (39), respectively.

These autoantibody profiling results help to outline the extent of broken B cell tolerance in our patient cohort and demonstrate patient to patient variability in autoantibody species and levels. It confirms and validates previously described APECED associated antibodies against cytokines with multiple assays, describes APECED associated autoantibodies against numerous enzymes agreeing with previous reports, and suggests the presence of many other autoantibody species that are less well understood prompting follow up studies (see supplemental spreadsheet). Next, to better understand how immune cell subsets might be changing in APECED compared to healthy individuals and potentially altering tolerance, we then turned to address how APECED impacts immune cell frequencies.

### Peripheral blood immune cells from APECED patients have a B cell compartment with reduced Naïve B cells, Transitional B cells, and increased Memory and Atypical B cell subsets relative to healthy donors

To explore how APECED affects the makeup of circulating immune cells, we stained patient peripheral blood cells for high parameter flow cytometry (see methods for details). We identified and analyzed the frequencies of major immune cell subsets including B cells, T cells, NK cells, and myeloid cells (Figure 2, A–I). We did not find changes in Tregs, TFHs, NKs, NKTs, and monocyte subsets in blood from APECED patients relative to healthy controls (Figure 2, A–I, and Supplemental Figure 2, A–I). However, we did observe alterations in B cell populations from the APECED patient samples (Figure 2). A significant reduction of the IgD+ CD19+ CD20+ B cell subset as a fraction of CD45+ cells was detected comparing Healthy (6.05 +/- 2.97%) and APECED (2.02 +/- 1.57 %) patients (Figure 2C). When looking within B cells, we found that there was a reduction in IgD+ B cells and increased phenotype of CD11C+ B cells and CXCR3+ B cells in APECED compared to healthy controls (Supplemental Figure 2, J– L). The fact that B cells are most altered in our APECED cohort is consistent with a recent longitudinal flow cytometry study showing B cells are most consistently changed in APECED across age (40).

**Figure 2.**
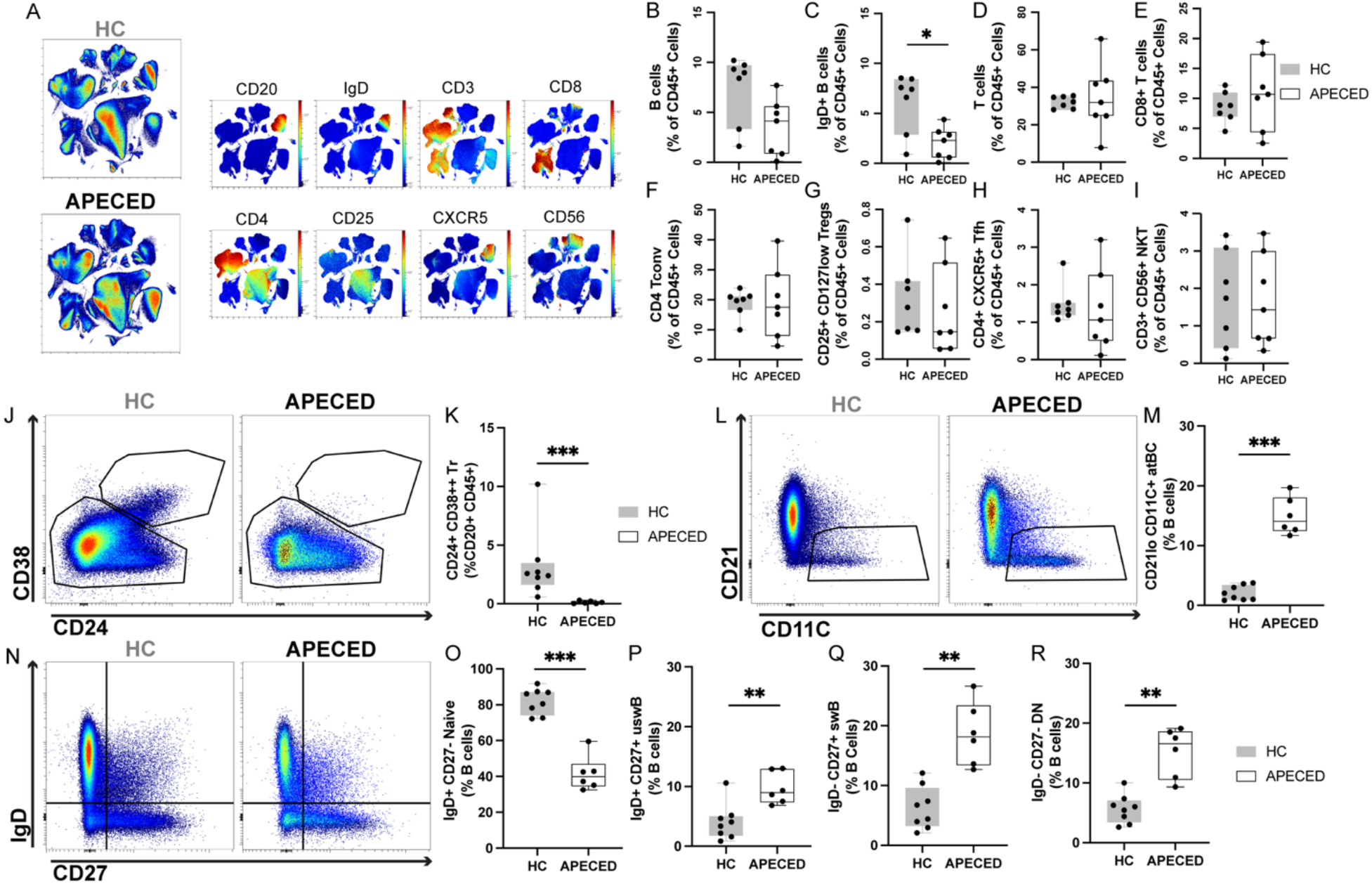
PBMCs from APECED patients have a B cell compartment with reduced Naïve B cells, Transitional B cells, and increased Memory and Atypical B cell subsets relative to healthy donors. (A) UMAP of live single CD45+ immune cells, concatenated for Healthy Control donors (HC) above and APECED donors below. MFIs for select immune lineage proteins within combined Healthy Control and APECED UMAP. (B-I) Quantification of immune cell frequencies shown as a percentage of CD45+ cells between Healthy Control and APECED. (J-R) B cell compositional analysis by flow cytometry. (J) Live single CD45+ CD20+ cells gated for transitional (Tr), and mature B cells in HC and APECED with quantification (J-K). Atypical B cell gating (L) and frequency quantification (M) comparing HC and APECED. (N) IgD/CD27 gating of mature B cells in HC and APECED with quantification of Naïve, uswB, swB, and DN subsets (O-R). Statistical tests for cell type percentages described in methods.

We further investigated the B cell compartment with a more focused flow cytometry panel in our cohort. We found significant reductions to Transitional B cells (Tr) as a proportion of CD45+ CD20+ cells in APECED patients (Figure 2, J and K). We uncovered substantial reductions in IgD+ CD27- Naïve B cells between Healthy (81.95 +/- 7.23%) and APECED (41.62 +/- 9.67%) patients, with increases in the proportions of CD21lo CD11C+ Atypical B cells (APECED 14.95 +/- 3.1%, HC 2.07 +/- 1.22 %), IgD+ CD27+ unswitched B cells (APECED 9.7 +/- 2.66 %, HC 4.07 +/- 3.04 %), IgD-CD27+ switched memory B cells (APECED 18.6 +/- 5.28 %, HC 6.26 +/- 3.58 %), and IgD- CD27- double negative DN cells (APECED 15.16 +/- 4.1 %, HC 5.64 +/- 2.39%) in APECED (Figure 2, L–R).

These changes demonstrate a significant shift away from naïve B cells and towards antigen experienced B cell subsets in APECED and prompt an in depth investigation into how APECED is impacting B cell phenotypes with single cell resolution.

### scRNAseq analysis confirms alterations to APECED B cell subsets

APECED B cells have not been profiled in molecular granularity with single cell methods. Our scRNA/BCRseq dataset provides a unique opportunity to understand the compositional changes and molecular signatures of immune subsets including B cells with isotype specificity. We first set out to map all sequenced peripheral blood cells from our patient cohort and understand how these subsets may be changed across conditions. We ran clustering, generated a UMAP plot, annotated immune cells by their top differential gene expression (Supplemental Figure 3, A and B, see supplemental spreadsheet for immune cell subset DEGs), and compared the frequencies of total peripheral blood cells between APECED and Healthy patients (Supplemental Figure 4, see methods for details). Further confirming our flow cytometry result, we observed a significant reduction in the overall percentage of Naïve B cells in APECED patient blood composition compared to Healthy control blood cells (Supplemental Figure 4A, Wilcoxon rank-sum Mann-Whitney tests, * p < 0.05). In addition to this observation, we saw a reduction in the frequencies of plasmablasts/plasma cells in APECED patients compared to Healthy (Supplemental Figure 4C). Although it did not reach significance, a potential trend was also seen towards higher CD14+ classical monocyte and intermediate monocyte populations in APECED compared to Healthy (Supplemental Figure 4, G and H). Differential gene expression analysis was carried out between APECED and Healthy conditions within each immune subset, results are available in a supplemental spreadsheet.

We then sought to follow up and understand the composition of the B cell compartment as a whole, taking all identified B cells from APECED and Healthy individuals for downstream clustering, UMAP, differential gene expression, and frequency analysis (Figure 3). We identified Naïve B cells (Naïve, IgD+ IgM+ TCL1A+ FCER2+ TBX3+ BACH2+), Activated B cells (aB, IgD+ IgM+ LINC01857+ ARHGAP24+ CTSH+) with downregulated expression of naïve B cell transcripts, Memory B cells (MBC, CD1C+ CD27+ TNFRSF13B+), switched memory B cells (swMBC, IgD- CD27+ LINC01781+ COCH+ TCF7+), atypical B cells (atBCs, FGR+ HCK+ ZEB2+ FCRL5+ ITGAX(CD11C)+ TBX21(Tbet)+ CD20++ CD19++ FGD4+ MPP6+), and plasmablasts/plasma cells (PB, CD27++ TNFRSF17 (BCMA)+ PRDM1(BLIMP-1)+, SLAMF7+ CD38++) (Figure 3, A and C).

**Figure 3.**
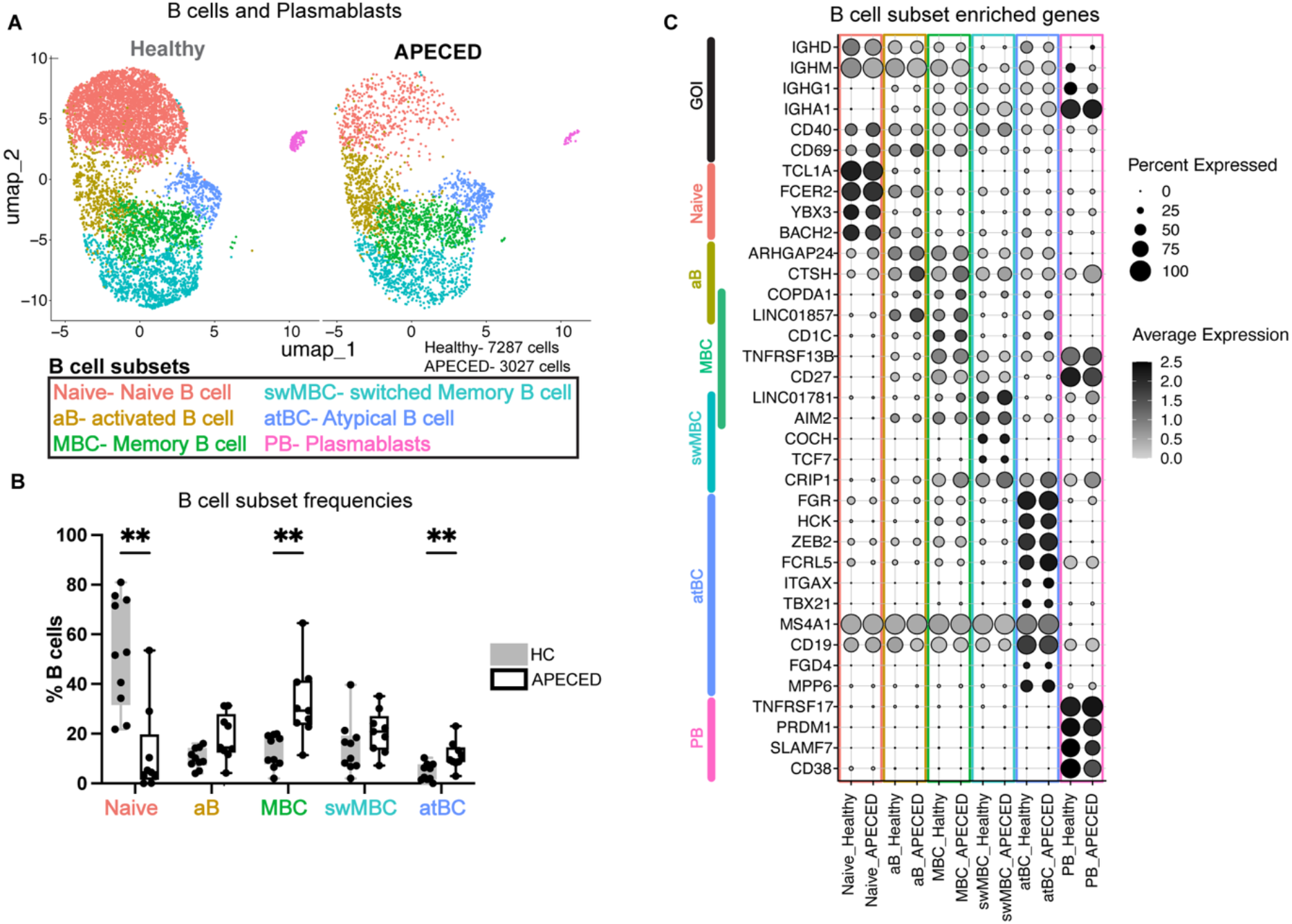
Changes in overall B cell subsets detected by scRNAseq. (A) UMAPs of B cells and PBs from Healthy and APECED conditions. 7287 Healthy and 3027 APECED cells. (B) Frequencies of B cell subsets in Healthy and APECED individuals. Wilcoxon rank-sum Mann-Whitney tests. ** p < 0.01 (C) Top differentially expressed genes in B cell subsets.

Overall we found substantial changes to the makeup of the B cell compartment in APECED patients, with a significant reduction in the frequencies of Naïve B cells, and significant increases to the frequencies of both MBCs and atBCs in APECED compared to Healthy (Figure 3B).

Due to the fact that both IgM and class switched IgG and IgA B cell populations contribute to MBC and atBC clusters that are being over-represented in APECED patients, it was unclear if further unique molecular subtypes were underlying these changes to APECED B cells. We then turned to address this question with the isotype specific resolution provided by BCRseq.

### B cell isotype specific subset clustering highlights compositional increases in alternatively activated IgM CD27+ B cells and class switched Atypical B cells

IgM and class switched IgA/IgG B cells were positively identified from sequenced peripheral blood cells with BCRseq, and then analyzed separately (see methods for details). Top differentially expressed genes were used to name B cell subsets, guided by learnings from B cell single cell RNAseq analysis (41) and naming conventions in agreement with recent efforts to align the field on B cell subset terminology (42).

In IgM B cells, we identified transitional (Tr, CD38+ SOX4+ MZB1+) and Naïve B cells (Naïve, IGHD+ TCL1A+ FCER2+ BACH2+) that were substantially reduced in samples from APECED patients (Figure 4, A, B, and D). These results confirm and supplement our flow cytometry results. Transitional B cells were nearly absent in our patient cohort, an observation that has been made in numerous previous reports, but whose cause is not yet known (11). We found significant increases in the frequencies of IgM CD27+ memory B cells (CD27+ MemB, CD27+ TACI+ MARCKS+ GPR183+) and IgM CD27+ CXCR3+ Atypical B cells (CD27+ atBCs, ZEB2+ TBX21+ ITGAX+ CRIP1+ CD27+ TACI+ EGR1+ CXCR3+ ANXA2+ CD1C+), collectively named IgM CD27+, while the proportion of Activated Naïve B cells (AcN, IGHD+ CD27- ZEB2+ TBX21+ ITGAX+ MPP6+ FGD4+) were not changed. (Figure 4, A, B, and D).

**Figure 4.**
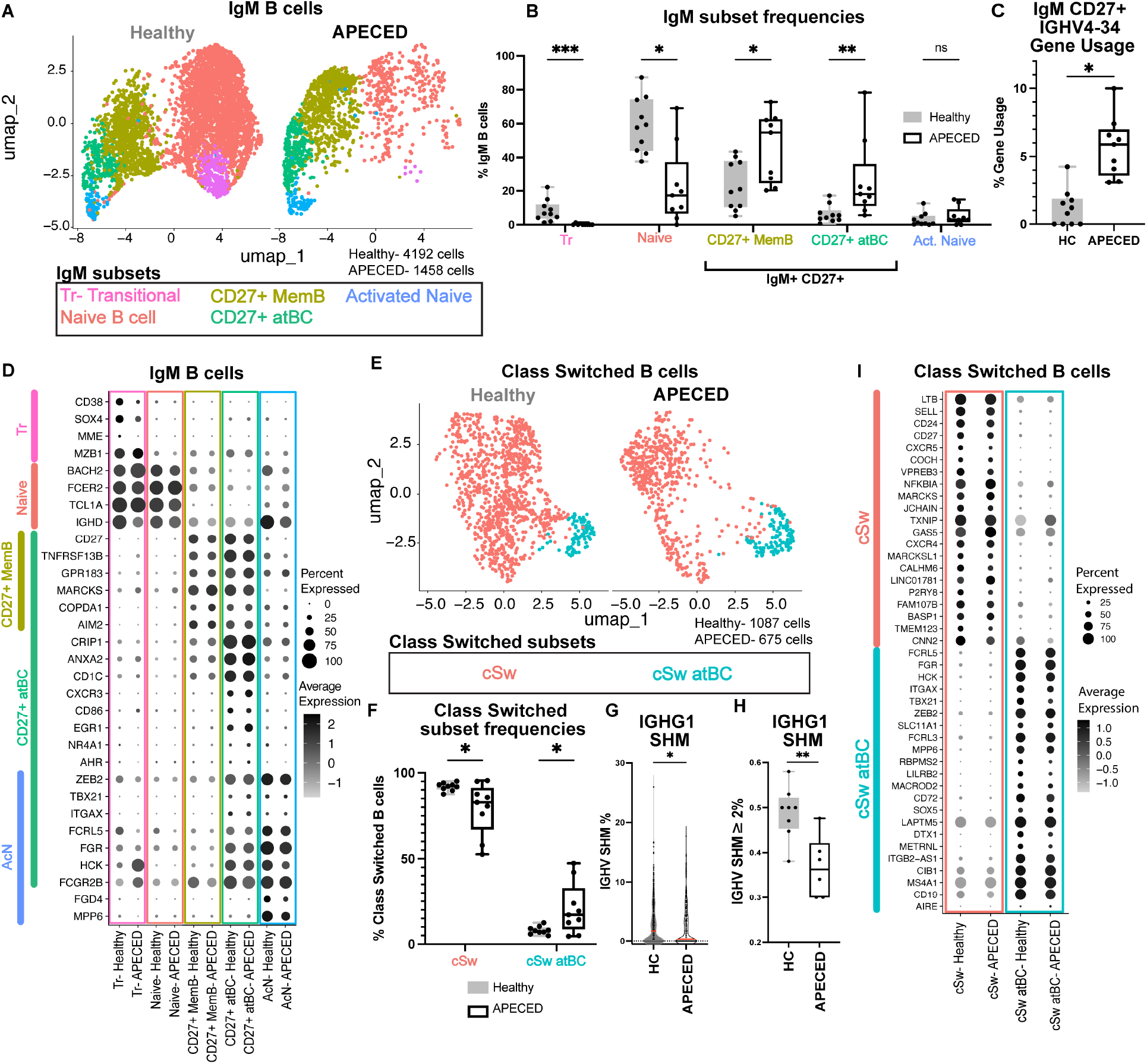
B cell isotype specific subset clustering highlights compositional increases in alternatively activated IgM CD27+ B cells and class switched atypical B cells. (A) UMAPs of IgM isotype B cells from Healthy and APECED patient blood. 4192 Healthy and 1458 APECED cells. (B) Frequencies of IgM subsets in Healthy and APECED individuals. IgM+ CD27+ B cells comprise CD27+ MemB and CD27+ atBC. Wilcoxon Rank sum Mann-Whitney tests. * p < 0.05, ** p < 0.01 *** p < 0.001. (C) Gene usage analysis of IGHV 4-34 in IgM+ CD27+ B cells. (D) Top differentially expressed genes in IgM B cell subsets. (E) UMAPs of class switched isotype B cells from Healthy and APECED patient blood. 1087 Healthy and 675 APECED cells. (F) Frequencies of class switched subsets in Healthy and APECED individuals. (G) BCR SHM was significantly lower in APECED IGHG1 B cells compared to healthy donor IGHG1 B cells in group comparisons. Wilcoxon Rank sum Mann-Whitney tests. (H) The proportion of SHM >2% in IGHG1 B cells per patient in APECED and Healthy donors (I) Top differentially expressed genes in class switched B cell subsets.

IgM CD27+ atBCs and Activated Naïve B cells together had a shared signature of 108 genes including ZEB2+ TBX21+ ITGAX+ FCRL5+ FGR+ HCK+ and FCGR2B+ (Figure 4D, and Isotype specific subsets Suppl DEG sheet), many of which have been described to support an alternatively activated B cell lineage that is seen in conditions of autoimmunity, infection, or chronic antigen exposure (42, 43, 44). It is noteworthy that the proportion of AcN does not increase in APECED unlike in SLE.

Interestingly, the expression of CXCR3 on atypical like B cells has been described as a phenotype seen in chronically stimulated B cells (42, 43); CXCR3 may also play a role in trafficking these B cells to inflamed organs through binding to its interferon gamma inducible chemokine ligands CXCL9/10/11. Unlike the CD27+ MemB cell subset, IgM CD27+ CXCR3+ Atypical B cells uniquely had an antigen presentation and anergic like gene signature as shown by the expression of CD86, NR4A1, AHR, and EGR1, similar to what has been previously reported in autoreactive prone and chronically stimulated B cell populations (45, 46).

We then evaluated the BCR repertoire between APECED and Healthy by performing IGHV gene usage analysis (see methods for details). Of all IGHV gene usage comparisons between APECED and Healthy B cells (IgM subsets, IGHG1, and IGHA1 B cells) we observed that only IGHV4-34 gene usage was significantly impacted in APECED IgM CD27+ B cells after correcting for multiple comparisons (Figure 4C). This result is particularly striking as germline IGHV gene IGHV4-34 is known to be autoreactive prone, binding to I/i red blood cell antigen, B cells, apoptotic cells, commensal bacteria, (47–51), and has been observed at a higher rate in SLE, eosinophilic granulomatosis with polyangiitis (EGPA), and Crohn’s disease (52).

These data show a major shift in the composition of IgM B cells away from Naïve B cells (Healthy average 59.2 +/- 16.2 standard deviation, APECED average 22.7 +/- 22.9 standard deviation) towards IgM CD27+ B cells (Healthy average 29.6 +/- 18.0 standard deviation, APECED average 64.7 +/- 31 standard deviation) that have a higher utilization of IGHV4-34 compared to Healthy IgM CD27+ B cells.

Class switched B cell subsets were identified in Healthy and APECED patients comprising IgG and IgA isotypes. Class switched memory B cells (cSw) and one subset of class switched atypical B cells (cSw atBC) were detected (Figure 4E) with top differentially expressed genes shown in Figure 4I (Isotype specific subsets Suppl DEG sheet). Class switched atypical B cells (cSw atBC) were found to be compositionally enriched in APECED patients compared to Healthy controls (Figure 4F).

cSw atBC expressed higher levels of genes characteristic of “DN2” like cells, classic alternatively activated atypical B cells (Eg. FCRL5, ZEB2, ITGAX, TBX21, FGR, HCK) (Figure 4I). These cells have been described in autoimmune conditions such as SLE (42, 43, 44) and are thought to be a potential source of autoreactivity.

We then asked how APECED may be affecting BCR phenotype through IGHV somatic hypermutation (SHM) analysis and found that only IGHG1 was significantly altered. Lower IGHV SHM rates were found in APECED IGHG1 B cells compared to Healthy IGHG1 B cells as a group (Figure 4G) and in the proportion of cells per patient (Figure 4H). Together these data show that the class switched B cell phenotype in APECED features a higher frequency of switched Atypical B cells and IGHG1 B cells with lower IGHV somatic hypermutation rates compared to Healthy controls.

We then evaluated differential gene expression across APECED and Healthy conditions in each isotype specific subset. The differential expression analysis comparisons are included in a supplemental spreadsheet and demonstrate increases to 20 genes shared across 2 or more subsets in APECED featuring mitochondrial oxidative phosphorylation and BCR isotype genes (discussed further in the results section below describing Supplemental Figure 6). They also feature unique upregulation and downregulation of transcripts in B cell subsets, especially in Naïve B cells with 21 unique upregulated genes, and Naïve, CD27 MemB, and cSw subsets showing unique downregulation of 13, 21, and 19 transcripts, respectively.

Overall, this highly granular analysis of B cells shows that APECED disease skews both IgM and class switched B cell subsets towards an alternatively activated phenotype, highlighting increases in the proportions of IgM CD27+ B cells (IgM Memory and IgM atypical B cells) as well as class switched atypical B cells.

### Proteomic analysis identifies substantial changes in serum biomarkers in APECED patients

To explore for potential drivers of APECED disease and factors that further characterize the APECED immune phenotype, we performed serum proteomics in our patient cohort (see methods for details). We found 334 proteins that were nominally different across the conditions, 267 upregulated and 67 downregulated in APECED patients (Supplemental Figure 5A). Two proteins reached experiment-wise significance: CXCL9 was increased and IL22 decreased relative to Healthy controls (Supplemental Figure 5A). An IFNG signature was evident through increases in serum CXCL9, CXCL10, IL12A/B, IL15, OX40, IFNGR1, and IFNGR2 (Supplemental Figure 5B). These results agree with previous reports that show APECED is driven by a Th1 response, where JAK inhibition has offered significant clinical benefit to patients (53).

In addition to the IFNG signature, there were nominal increases in acute phase response proteins (SERPINA1, LBP, HMOX2, C1QA, C5, CFP), innate immune system (REAC:R-HSA-168249, p = 0.00745, Eg. CD14, PGLYRP1, TREM2, FCN1, CLEC5A, RNASE6) and especially myeloid associated proteins including myeloid leukocyte migration (GO:0097529, p = 1.78E-07, Eg. CCL3, CCL14, CCL16), and proteins related to tissue damage and repair (TGFBR1, TGFBR2, SPP1, CSF1, TNFRSF1A, TNFRSF1B), among others (OLINK Suppl spreadsheet all comparisons).

We then questioned whether immune cells were being influenced by IFNG using a IFNG signature gene score (Supplemental Figure 5, C and D) and found that APECED immune subsets had a significantly increased IFNG gene score including T cells, myeloid cells, and B cells (Supplemental Figure 5C). This supports the notion that an IFNG response is prevalent in APECED patients as previously described.

### Functional AIRE expression does not significantly impact the transcriptomic state of class switched atypical B cells detected by scRNAseq

It has been observed that AIRE has a cryptic role in human atypical B cells that may influence tolerance (54). To help address the question on how functional AIRE expression influences the classic atypical B cell signature, we looked for the presence of AIRE expression in our B cell dataset and found that AIRE RNA is indeed detected in 5-10% of class switched Atypical B cell populations in both Healthy and APECED patients (Supplemental Figure 6A). However, a differential expression analysis between APECED and Healthy class switched Atypical B cells yielded only 9 genes (log2fold = 0.6, min pct = 0.1). All 9 genes were mitochondrial oxidative phosphorylation genes upregulated in APECED (GO:BP Oxidative phosphorylation, p = 7.508 x 10^-16) (Supplemental Figure 6B). Importantly, these changes to mitochondrial oxidative phosphorylation genes were also upregulated in APECED compared to Healthy in other subsets such as IgM CD27 atBCs (8/9 upregulated Oxidative phosphorylation genes shared), class switched memory B cells (7/9 upregulated Oxidative phosphorylation genes shared), and IgM MemB (5/9 upregulated Oxidative phosphorylation genes shared) which show a group mediated effect rather than a subset specific effect. We then examined the Aire regulated Tissue Restricted Antigen (TRA) gene score metric from previous studies (54, 55) in our cells and found no significant differences in TRA gene scores between cSw atypical B cells with functional (Healthy) or mutated (APECED) copies of the AIRE gene (Supplemental Figure 6, C and D). While the limitations of 10X genomics capturing only a fraction of the single cell transcriptome in individual cells makes it impossible to completely rule out a direct effect of AIRE expression in B cells, these data do not support a role for AIRE in regulating gene expression in classic atypical B cells.

### cSw Atypical B cells are prone to autoreactivity irrespective of AIRE deficiency

Since class switched atypical B cells (cSw atBCs) are enriched in APECED and have been previously tied to autoantibody production in autoimmune conditions such as SLE, we wondered whether this subset could be a potential source of autoreactivity. To test this, we cloned 48 antibodies derived from APECED cSw atBCs and 37 from Healthy cSw atBCs (Figure 5, A and B) from our BCRseq dataset along with 176 random healthy donor IgG+ B cells and screened them for the binding of autoantigens in two separate antigen panels (see methods for details). In our selection, we sampled cSw atBCs BCRs with IgG (APECED = 30, Healthy = 27) and IgA (APECED = 18, Healthy = 10) isotypes.

**Figure 5.**
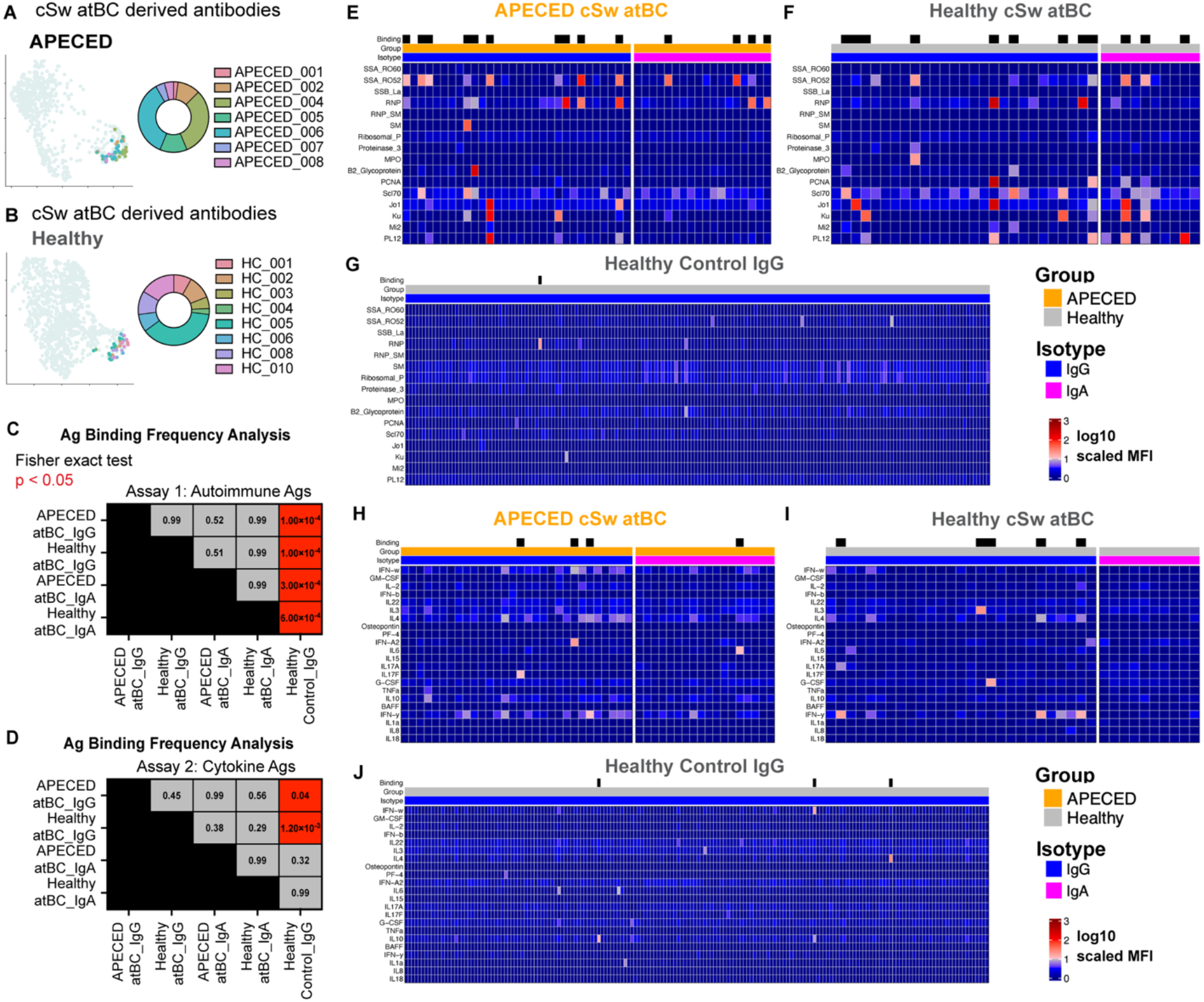
Class switched Atypical B cells are autoreactive prone irrespective of AIRE deficiency. (A-B) (A) APECED and (B) Healthy class switched atypical B cell derived antibodies. Patients represented on right, and placement in scRNAseq class switched UMAP on left. 176 IgG+ B cell antibodies from healthy donors were used as a control. (C) Assay 1 autoimmune antigen binding frequency analysis for APECED atBCs and Healthy atBCs compared to Control IgG+ B cell derived antibodies from panels E-G data. Fisher exact test p values for each comparison. (D) Assay 2 cytokine antigen binding frequency analysis for APECED atBCs and Healthy atBCs compared to Control IgG+ B cell derived antibodies from panels H-J data. Fisher exact test p values for each comparison. (E-G) Antibodies were screened for binding with Assay 1 (E, F, G) and Assay 2 (H, I, J) autoantigens. Each column is an antibody and each row is an antigen, arranged by group and isotype tested. APECED groups labeled in orange and Healthy in grey. IgG isotypes labeled in blue and IgA labeled in pink. Binding antibodies are labeled on the top row with a black bar.

When we assayed the cloned cSw atBC antibodies against a panel of autoimmune disease antigens (SSB/RO-60, SSA/RO-52, RNP, MPO, Mi2b, Sm/RNP, Sm, Scl70, Jo1, SSB/La, B2 Glycoprotein, PR3, PCNA, Ribosomal P, PL12, Ku) we found that 10/30 of the APECED IgG atBC antibodies and 4/18 APECED IgA atBC antibodies met the criteria for binding one or more antigens (Figure 5E) compared to 1/176 control IgG antibodies (Figure 5G). Both APECED IgG and IgA atBCs had a higher proportion of antibodies binding autoantigens than control IgG B cells as determined by fisher exact test (Figure 5C). Similarly, when we compared cSw atBCs derived antibodies from healthy patients we saw an increased proportion of autoantigen binding antibodies in IgG (9/27 antibodies) and in IgA (3/10 antibodies) (Figure 5F) compared to Healthy control IgG antibodies (Figure 5, G and C). However; when APECED (Figure 5E) or healthy (Figure 5F) atypical B cell antibody binding frequencies were compared, there were no significant changes (Figure 5C). Therefore, cSw atBCs of IgG and IgA isotypes from either APECED or Healthy patients are more likely to bind these autoimmune antigens compared to control IgG B cells.

We then tested these antibodies against a panel of cytokine antigens (IFN-w, GM-CSF, IL-2, IFN-b, IL22, IL3, IL4, Osteopontin, PF-4, IFN-A2, IL6, IL15, IL17A, IL17F, G-CSF, TNFa, IL10, BAFF, IFN-y, IL1a, IL8, and IL18) that included autoantigens found in APECED patients (Eg. IFN-w, IFN-A2, IL17F, IL17A, IL22, IL6) and found that 3/30 APECED IgG atBC antibodies bound antigens (IFN-A2, IL17F, and IFN-y, Figure 5H) and 1/18 APECED IgA bound antigen (IL6, Figure 5H) compared to 3/176 control IgG antibodies that bound antigen (IL10, IFN-w, IL4, Figure 5J). APECED IgG atBCs but not APECED IgA atBCs had a higher proportion of binding antibodies to cytokine antigens compared to Control IgG+ B cells (Figure 5D). Similarly, Healthy IgG atBCs (5/27 antibodies, Figure 5I) but not IgA atBCs (0/10 antibodies, Figure 5I) had a higher rate of binding cytokine antigens than control IgG antibodies (Figure 5, J and D). This result shows that IgG cSw Atypical B cell derived antibodies sourced from either APECED or Healthy donors were more likely to bind cytokine autoantigens than healthy IgG control B cells (Figure 5D), tested by fisher exact tests (APECED IgG atBC vs IgG Control: p value = 0.04; Healthy IgG atBC vs IgG Control: p value = 0.0012).

These assays together show cSw atBCs as a cell type are more prone to bind autoantigens than IgG control antibodies. The over-representation of cSw atBCs in the APECED B cell compartment may therefore help contribute to the breakdown of peripheral B cell tolerance.

## DISCUSSION

Our immune profiling analysis from the peripheral blood of APECED patients reveals diverse serum autoantibody profiles and notable alterations in B cell composition. We observed a reduction in transitional and naïve B cells, coupled with an increase in alternatively activated populations including IgM CD27+ and class-switched atypical B cells whose BCRs exhibit autoreactive prone features.

IgM CD27+ B cells were comprised of IgM memory (IgM MemB) and IgM atypical B cell (IgM atBC) phenotypes that were both expanded in the APECED B cell compartment compared to Healthy control IgM B cells. IgM CD27+ atBCs had overlapping signatures with both IgM CD27+ MemB (Eg. shared IgDlo CD27+ TNFRSF13B+ MARCKS+ expression) and Activated Naïve (AcN) subsets (Eg. shared Tbet+ ZEB2+ FCRL5+ HCK+ FGR+ expression) as well a as unique expression profile characterized by CXCR3 chemokine receptor, CD1C, CD86, and markers for anergy such as AHR, EGR1, and NR4A1. This combination of markers suggests that IgM CD27 atBCs may be chronically activated. The expression of Tbet and CXCR3 on IgM CD27 atBCs also supports the notion that this subtype may be driven by IFNG.

BCR analysis showed that APECED IgM CD27+ B cells frequently utilize the IGHV4-34 variable chain gene (also known at the 9G4 idiotype), which has been previously seen expanded in autoimmune diseases such as SLE, EGPA, and Crohn’s disease (52) as well as cold agglutinin disease and numerous cancers (50, 51). The germline IGHV4-34 FR1 hydrophobic patch region necessary for the anti- Ii red blood cell antigen and anti-B cell activity of the 9G4 idiotype was found intact in APECED IgM CD27+ B cells. There is recent attention in using this target to treat autoimmunity and cancer indications by depleting this B cell subset with bispecific and especially CAR-T approaches (56, 57).

Compositional analysis of class switched B cells showed that class switched atypical B cells (“DN2”) are expanded in APECED patients compared to healthy controls. Class switched atypical B cells have been observed increased in autoimmune conditions (eg. SLE, Rheumatoid arthritis, Sjogrens syndrome), infections (eg. HIV, Malaria), and during numerous LOF innate errors of immunity affecting genes such as CTLA4, LRBA, FAS, STAT5B, and PDCD1 (58). Exactly why these changes are occurring in APECED is not clear, but there may be a convergent role for T cells as the aforementioned LOF innate errors of immunity that affect T cell activation and tolerance drive increases in atypical B cells. Additionally, there has been a proposed role for systemic inflammation and interferon gamma which was observed in the serum of our APECED cohort that has been suggested to help drive this phenotype.

Interestingly, the proportion of AcN B cells were not significantly altered in APECED. This AcN population has been observed increased in SLE and is thought of as a precursor to the class switched atypical “DN2” phenotype. Further study is needed but these differences could suggest different routes of B cell activation between APECED and SLE that may converge on a phenotypically similar class switched atypical “DN2”.

Since the class switched atypical “DN2” like B cell subset has been observed in autoimmune conditions and is thought to be autoreactive prone, we cloned antibodies from the sequences of these class switched atypical B cells of APECED and Healthy patients and tested them in autoantigen binding assays, comparing them against IgG B cell controls. Class switched atypical B cells from both healthy and APECED demonstrated a higher propensity for autoantigen binding compared to control IgG B cells, especially cSw atBCs of the IgG isotype. The APECED cSw atBCs tested were most likely to bind RNP and RO52 in this assay, and included a number of antibodies that bound to multiple antigens. Very few cSw IgG atBCs bound to cytokine antigens, but of the three APECED cSw atBCs that met the binding criteria of the screen, all three of them bound to the APECED associated antigens IFN-A2, IL17F, and IL6.

However, healthy and APECED patients’ switched Atypical B cells were similar in their likeliness to bind panel autoantigens overall and despite varying in functional and mutant AIRE, had very few differentially expressed genes between them including a similar expression of AIRE regulated peripheral tissue antigens as measured by their peripheral tissue antigen gene score with scRNAseq. It is worthwhile noting however, that this data is limited by the lower sensitivity and sequencing depth of single cell methods. Nonetheless, this offers a perspective in which cell state may be the determining factor toward atypical B cell autoreactivity regardless of Aire deficiency. The increased proportion of these cells in APECED therefore may contribute to the breakdown of peripheral B cell tolerance. Furthermore, IGHG1 IGHV somatic hypermutation levels were found to be lower in APECED, suggesting that affinity maturation may be impaired in APECED patients. This evidence builds on a previous effort to understand how APECED skewing of the T cell repertoire impacts germinal center reactions and the potential dysregulation of lymphoid structures including the makeup of T follicular helper cells and Tregs (31).

Interestingly, the shifting composition of the B cell compartment was observed more in APECED patients that had a higher number of manifestations than those with low numbers of manifestations in this cohort. It may be worthwhile for future studies to track these changing B cell subsets in larger cohorts of APECED patients to correlate with disease features and therapeutic responses to see if they have biomarker potential.

Previous reports have observed changes in APECED B cells including increases to IgM CD27+ B cells (59) and those with an atypical like phenotype (12). Our study now provides the added molecular granularity of combined BCR and single cell transcriptomic analysis, highlighting that these isotype specific subsets have alternative activation signatures and feature autoreactive prone BCRs. Additionally, serum proteomics from our study and others indicate APECED patients’ pro-inflammatory environment related to IFNG and innate immune activation. This inflammatory milieu may help drive the observed phenotypic changes to immune cells including, among other effects, promoting an environment that could favor B cell activation.

Altogether, our findings support the notion that, along with the reported increase in autoreactivity in mature naïve B cells (9) due to a faulty peripheral tolerance checkpoint on B cells by APECED T cells, B cell activation in APECED causes a compositional shift towards different autoreactive prone IgM CD27+ and class switched atypical B cell subsets that may contribute to the conditions of broken B cell tolerance, subsequent autoantibody production, and autoimmunity.

## MATERIALS AND METHODS

### Sex as a biological variable

Our study cohort examined both male and female APECED and healthy subjects (Supplemental Table 1). Each group contained 10 individuals that were age and sex matched to control for these variables, totaling 7 females and 3 males per group.

### APECED study collection & sample processing

Whole blood and serum were collected from patients diagnosed with APECED (N = 10) and from age and sex matched controls (N = 10) by Sanguine Biosciences, Inc (Waltham, MA) If available, age matched healthy siblings were collected. For detailed patient information see Supplemental Tables 1 and 2.

Samples were shipped overnight and processed immediately upon arrival to isolate peripheral blood mononuclear cells (PBMCs) using SepMate tubes (STEMCELL, Ref # 85450). Blood was diluted 1:1 with PBS then added to SepMate-50 tubes filled with 15 mL of Ficoll-Paque PLUS (Cytiva, #17144003) and centrifuged at 1,200 g for 10 minutes. PBMCs and plasma layers were poured in a new tube, pelleted at 300 g for 10 minutes, and resuspended with 5 mL of ACK lysing buffer (Gibco, # A10492-01) for 3 minutes to eliminate red blood cells. PBS was added to 50 mL and PBMCs were pelleted at 300 g for 10 minutes. If necessary, the red blood cell lysis step was repeated. Portions of resuspended PBMCs were then used for single cell OMICs (scRNA/BCRseq) and high parameter flow cytometry. The remaining PMBCs were frozen in cryovials. All blood samples were handled in a BSL-2 laboratory with personal protective equipment and safety precautions, in accordance with the blood processing protocol approved by the Regeneron Institutional Biosafety Committee.

### Flow cytometry

PBMCs were blocked with Human TruStain FcX and True-Stain Monocyte Blocker (Biolegend) and stained with a 39 parameter immunophenotyping panel in 96 well U bottom plates. For each patient, 3 million cells were stained, and the entire sample was acquired in tubes with a Cytek Aurora. The flow cytometry antibodies and complete protocol for this panel are listed in a supplemental spreadsheet.

### scRNA-seq pre-processing

CellRanger (v.6.1.1, 10X Genomics) with GRCh38.100 reference was performed on the processed cells to align the reads, to calculate gene expression levels, and to make raw count matrix, gene/barcode list. Quality control has been done on the data to filter out abnormalities, empty droplets, and multiplets. In each sample, cells with the total number of molecules detected within a cell higher than 20,000 and lower than 2,000, and the total number of genes per cell higher than 4500 and lower than 800 were filtered out. Then Python module Scanpy (v1.7.1) was used to normalize the data. There were 3 steps for the normalization: 1. For every cell, scale total counts sum up to 10,000. 2. Log-normalization with log1p function. 3.Unit scaling the counts between 0 and 10.

R Seurat package (v5.0.1) was utilized to generate UMAP plots and to run clustering. In each cluster, we identified markers based on differentially expressed genes, and cell types were given to the clusters based on the top markers. For detailed annotation of cell types, we sub-clustered the cells separately for the B cells and T cells and compared those sub-clusters across in each cell type. Then detailed cell type annotations were given to each sub-cluster based on their top markers.

### scBCR-seq pre-processing and V(D)J analysis

Single-cell BCR and data were processed using the same version of CellRanger with the corresponding GRCh38.100 V(D)J reference. All included gene expression and V(D)J reactions passed CellRanger quality control metrics.

### OLINK explore & serum proteomics

Serum samples were processed using all 8 panels in the Olink Explore 3072 kit (Cardiometabolic I/II, Inflammation I/II, Neurology I/II, and Oncology I/II) using a custom workflow developed at the Regeneron Genetics Center and approved by Olink. The samples were sequenced on the Illumina NovaSeq 6000 platform on S4 flow cells according to Olink’s custom sequencing recipe. Plate normalized data was analyzed using the OlinkAnalyze library in R using the olink_ttest function (Welch’s 2 sample t test, alternative = “two.sided”, with Benjamini-Hochberg method correction using stats::p.adjust).

### Autoantibody profiling analysis

Serum autoantibody profiling was run according to the manufacturer’s instructions (HuProt, CDI laboratories). After blocking with 5% BSA/1xTBST at room temperature for 1 hour, the arrays were probed with the serum samples (1:500) at room temperature for 1 hour and washed with TBST for 10 min X 3, and then probed with Alexa 647-human IgG Fc gamma specific secondary antibodies under conditions optimized by CDI Labs for signal detection. Data was quantile normalized to control for inter-sample variability and T tests were performed comparing groups to explore for potential autoantibody varieties. Antigens increased 2 fold or greater in average MFI values in APECED over healthy and whose p values < 0.05 were labeled putative APECED associated autoantibodies.

### Generation of B cell derived antibodies and IgG ELISA

Antibodies were generated using heavy and light chain antibody variable regions sequenced from class switched atypical B cells of APECED and Healthy donors. These regions were cloned as human IgG4 and expressed as recombinant antibodies in Expi293F cells. Control IgG B cells were isolated from frozen PBMCs of two healthy donors (STEMCELL Technologies), sorted into 384-well plates, and their heavy and light chain antibody variable regions amplified by PCR. The regions were cloned as human IgG1 and expressed as recombinant antibodies in CHO cells. IgG ELISAs were then carried out to determine the concentrations of each antibody supernatant using human IgG standard (Jackson 009-000-003). Plates were coated overnight with 1:15K (0.1 uG/mL) F(ab)2 fragment anti F(ab)2 IgG (Jackson 109-066-097 lot 165925). Plates were washed 3x and then blocked in 1% BSA in PBS for 1 hour. Plates were washed 3x, samples were added at 1:10 dilution along with standard (from 5000 ng/mL to 6.86 ng/mL, 3 fold dilutions) and incubated for 1 hour. After washing 3x, detection antibody was used at 1:10,000 (Jackson 109-035-098) and incubated for 1 hour. After 3 washes, TMB was added for 15 minutes followed by stop solution and reading at 450 nm.

### Autoantigen assays

Patient serum and B cell derived antibodies were tested against two panels of antigens, the Milliplex Human Cytokine Autoantibody panel (HCYTABG-17K) and Milliplex Human Autoimmune Autoantibody panel (HAIAB-10K). Protocols were carried out according to the manufacturer’s instructions and read on a Luminex FLEXMAP 3D instrument. Positive control autoimmune serum SSB-La-HSB-0100, RNP-HRN101, SCL70-HSC-0100, Jo1-HJO-0100, SM-HSM-0100, (immunovision) and antibodies against PCNA (BD Biosciences, mouse monoclonal IgG1, clone 24/PCNA, cat # 610664) and RO-52 (Santa Cruz, mouse monoclonal IgG1, clone D12, cat # sc25351) were used to independently validate the presence of antigen. Control antibodies were tested in dilutions ranging from 4 ug/mL to 0.004 nG/mL. We found that specific binding signals with >10 fold ((Target Signal) / (median MFI for all beads per antibody)) could be detected from 4 ug/mL to 4nG/mL. Screened antibody supes were added in this concentration range. For cytokine antigens, APECED serum validated the presence of IFN-A2, IFN-w, IL17A, IL17F and IL22. Binding criteria were met when the antibody MFI signal detected in one or more antigen bead was both (1.) 10 fold + 5x StDev or greater than supe background control per the antigen; and (2.) 10 fold or greater than the median MFI for all antigens per antibody. This helped assure that the detection was both high signal over background noise and specific.

### Statistical Analyses

Differential expression analyses were performed using the nonparametric Wilcoxon rank-sum test with the Seurat package (version 5.0.1) in R (version 4.2.1) using the FindMarkers function (min.pct = 0.1 and logfc.threshold = 0.6). These comparisons included global (each cluster versus all others) and pairwise differential expression analyses across clusters for functional annotation. We corrected the P values for multiple testing with the Benjamini-Hochberg procedure, and considered genes with an adjusted P < 0.05 as statistically significant.

Pathway analysis was done to up-regulated or down-regulated genes of the comparisons. GSEA (Gene Set Enrichment Analysis) on mSigDB, clusterProfiler package (version 4.6.2) with GO (Gene Ontology) and KEGG database in R, and g:Profiler (https://biit.cs.ut.ee/gprofiler/gost) were used for the pathway analysis to provide associated biological functions to the genes of interest.

Cell type percentages were calculated by [the number of cells for the cell type x 100 / the total number of cells in the condition]. Distributions were assessed for normality with Shapiro-Wilk tests. Unpaired T tests were used for data that passed this normality test and Wilcoxon rank-sum tests were used on distributions that failed to pass normality. Percentages of IGHV gene usage were calculated by [the number of cells with the IGHV gene x 100 / the total number of cells in the condition]. Comparisons for the percentage of IGHV gene usage within B cell conditions were performed using the non-parametric Wilcoxon rank-sum test using the R package rstatix. P values were then corrected with the Benjamini Hochberg procedure, and IGHV genes with adjusted P < 0.05 were considered significant between groups.

### Study approval

Human subject samples were obtained from Sanguine Bioscience (Waltham, MA) following protocol approval (Protocol: Prospective Collection of Whole Blood from Healthy Patients and Patients Diagnosed with Autoimmune polyendocrinopathy-candidiasis-ectodermal dystrophy (APECED), Protocol Number SAN-09166) from the Institutional Review Board (IRB). Written informed consent was received prior to participation.

## Supporting information

IsotypeBcells_Subsetmarkers

IsotypeBcells_GroupDEGenes

PanB_SubsetMarkers

PanB_GroupDEGenes

PeripheralBlood_Subsetmarkers

PeripheralBlood_GroupDEGenes

HuProtArray

Olink_NPXvalues

FlowCytometry_Panels

## Data availability

Single cell multi-omics data and proteomics data is submitted to GEO (GSE339254) and will be made publicly available. Flow cytometry, autoantibody profiling, differential gene expression analyses, and assay data is provided in supplemental material. Any additional data and material from the study will be available upon request.

## Author contributions

Conceived and designed studies: A.A.B., S.H., and W.G. Methodology: A.A.B., S.H., W.G., C.A., H.K., B.K., and M.M. Conducted experiments: W.G., B.K., and M.M. Acquired data: W.G., B.K., and M.M. Analyzed data: W.G., H.K. Wrote manuscript: W.G. Reviewed manuscript: All authors.

## Acknowledgments.

We are grateful to the patients and their families for their participation in our study. We would like to thank Brace Porter, Christina Yip, Feiin Lou, Isabel Kao, Natalia Swiecki, Bei Wang, Jacquelynn Golubov, John Overton, Sean O’Keeffe, Hang Du, Erin Brian, and Kristy Guevara for their technical support and assistance. W.G. thanks the Regeneron Postdoc program and the Postdoc Steering Committee for their support and helpful feedback. We thank Seblewongel Asrat, Andrea Vecchione, Eva Conde Garcia, Valerio Donato, Erica Ma, Chen Shu Dong, and Angelos Papatheodorou for their helpful discussions.

---

**Supplemental Figure 1 S1.**
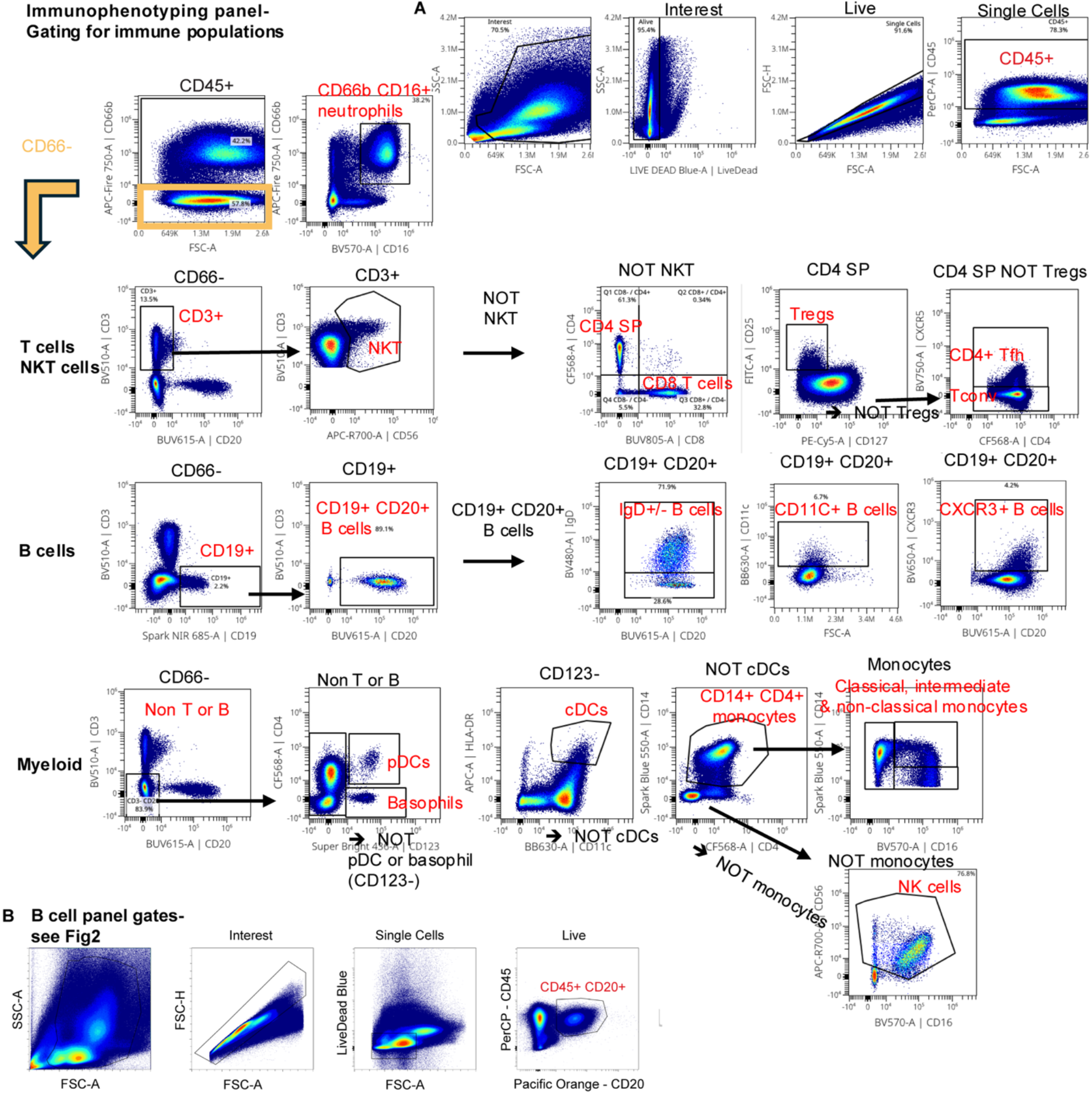
(A) Gating strategy for peripheral blood immunophenotyping. (B) Gating strategy for B cell panel.

**Supplemental Figure 2 S2.**
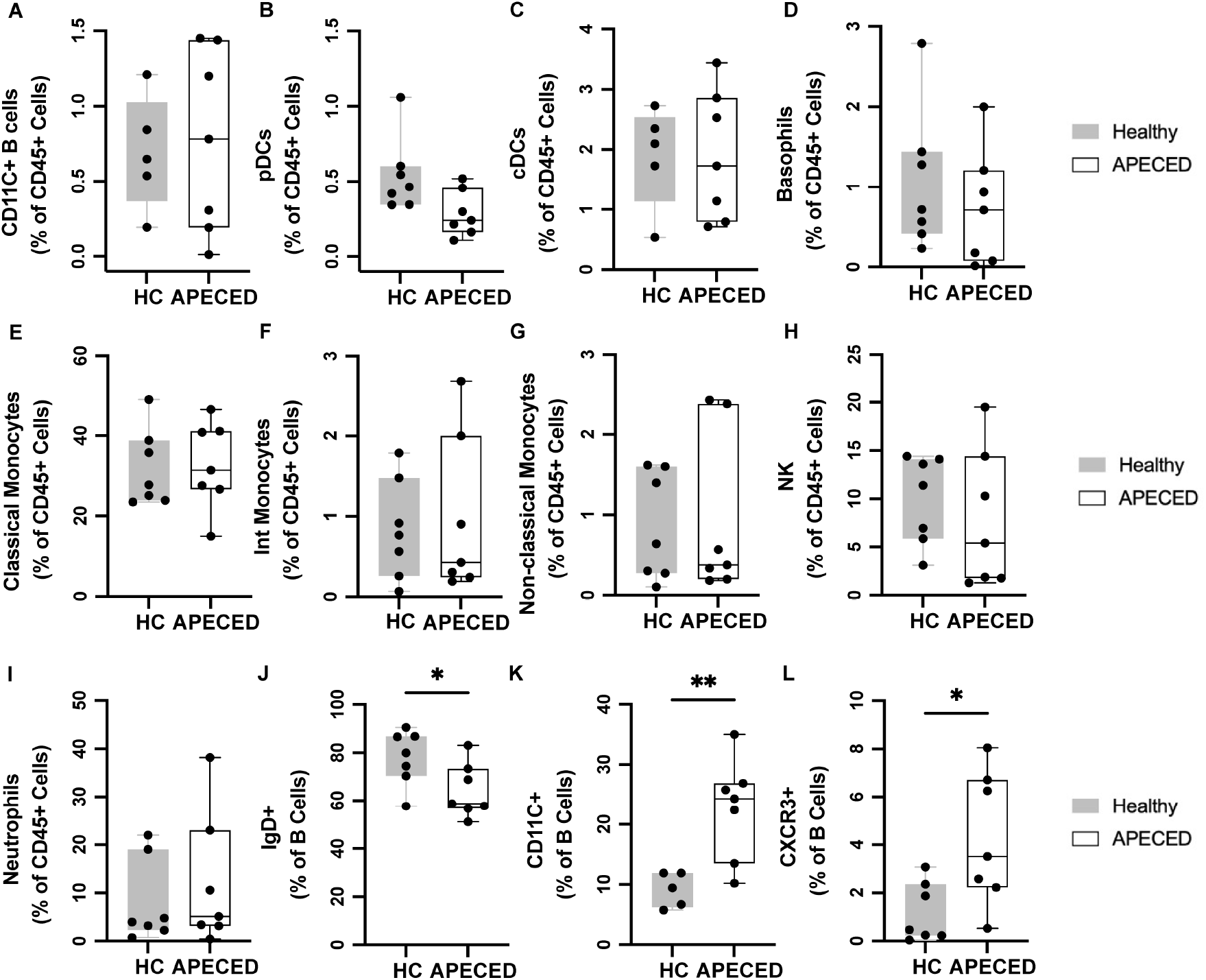
Immune frequencies from peripheral blood flow cytometry immunophenotyping comparing Healthy and APECED patients continued. **(A-I)** Quantification of immune cell frequencies shown as a percentage of CD45+ cells between Healthy Control and APECED continued from Fig 2. (J-L) Quantification of subsets as a percentage of CD19+ CD20+ B cells. Two tailed Wilcoxon rank-sum test. * p < 0.05. ** p < 0.01.

**Supplemental Figure 3 S3.**
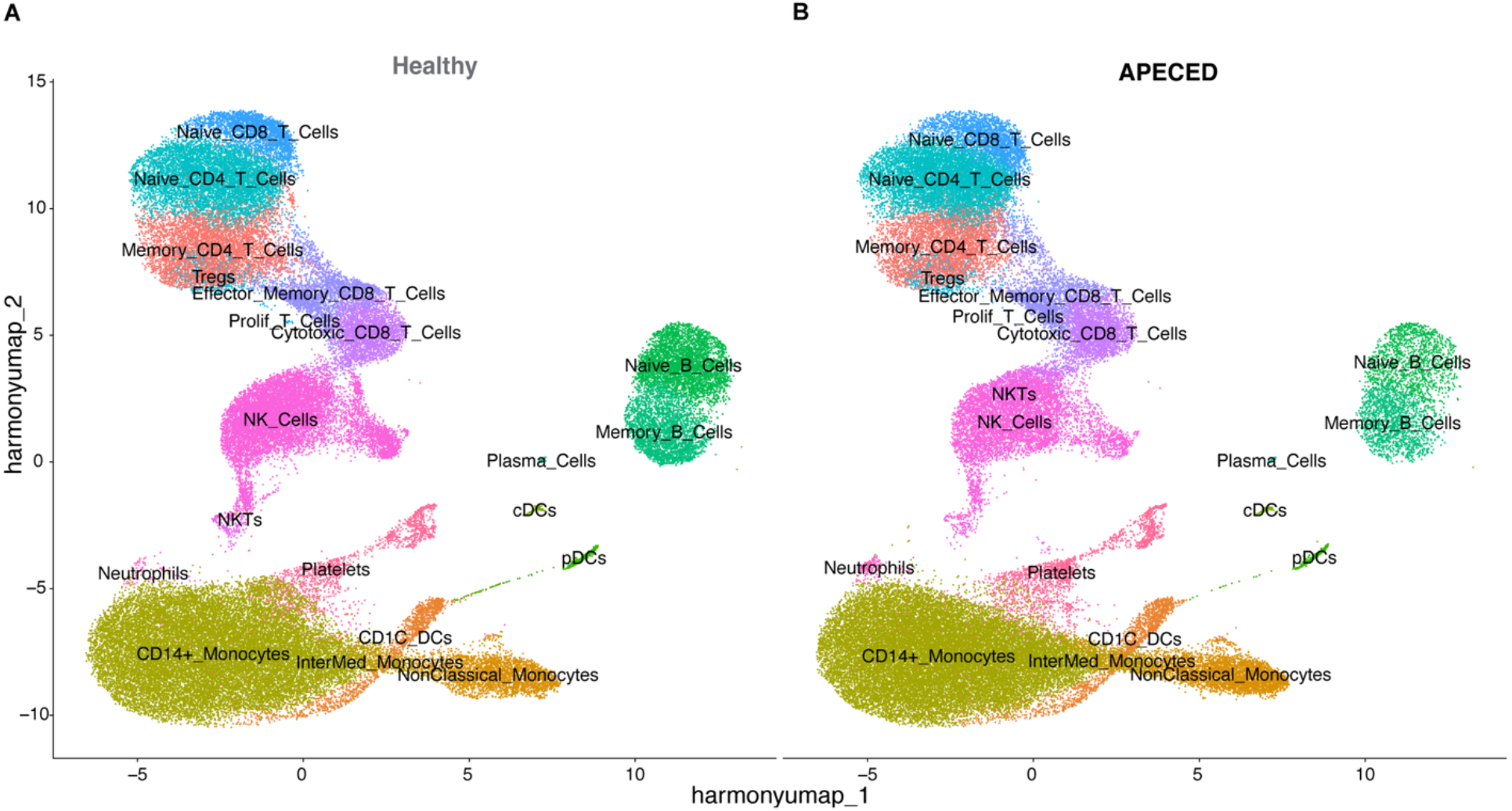
UMAP and frequencies of all peripheral blood cells sequenced. (A, B) UMAP of all peripheral blood cells sequenced from Healthy (left, A) and APECED (right, B) donors. Immune subsets labeled. Representing 67154 cells from Healthy patients and 64102 cells from APECED patients.

**Supplemental Figure 4 S4.**
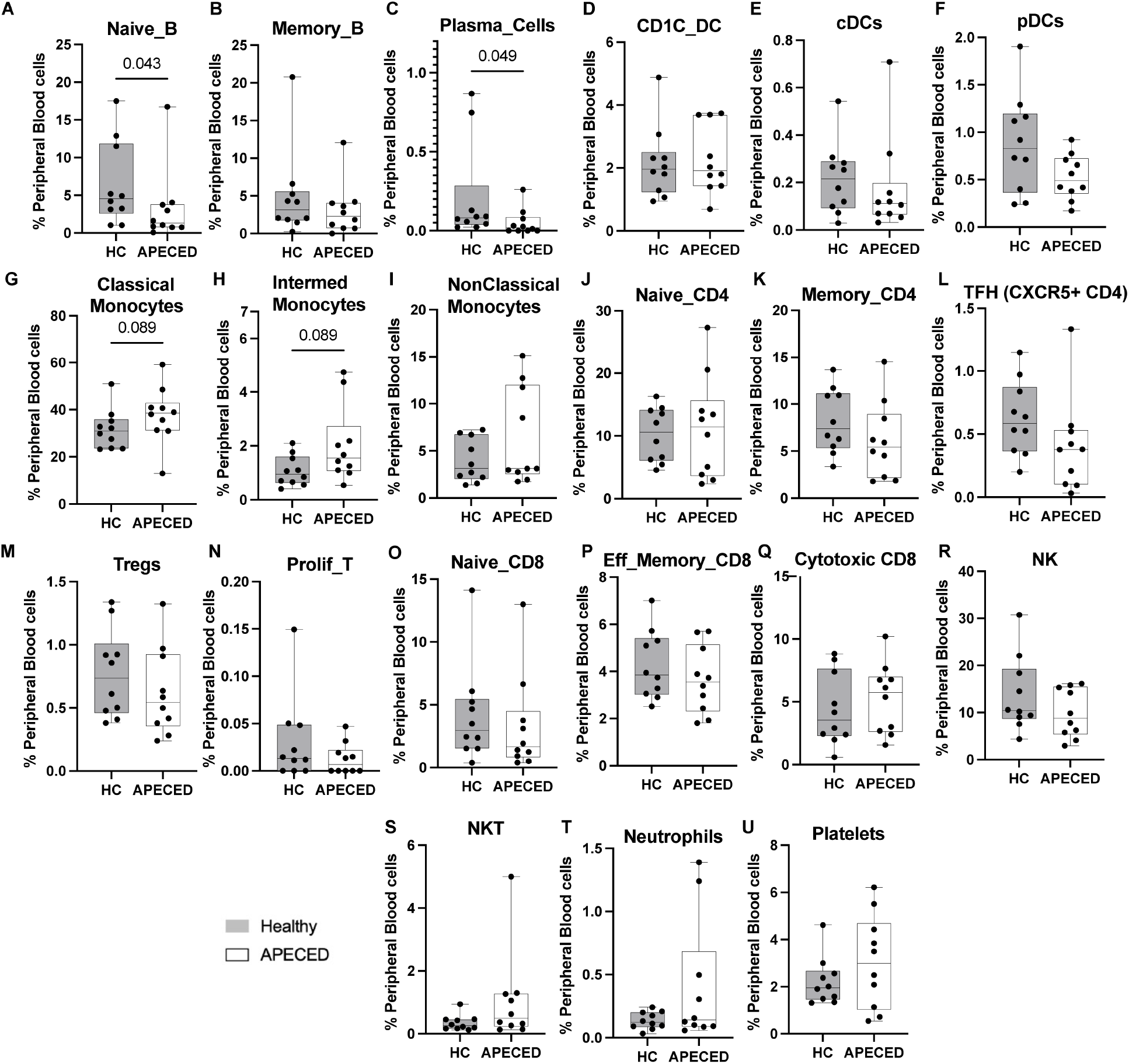
Immune subset frequency analysis from all sequenced peripheral blood cells comparing Healthy and APECED patients. **(A-U)** Quantification of immune subset frequencies shown as a percentage of peripheral blood cells between Healthy Control and APECED. Two tailed Wilcoxon rank-sum test. P values < 0.1 shown.

**Supplemental Figure 5 S5.**
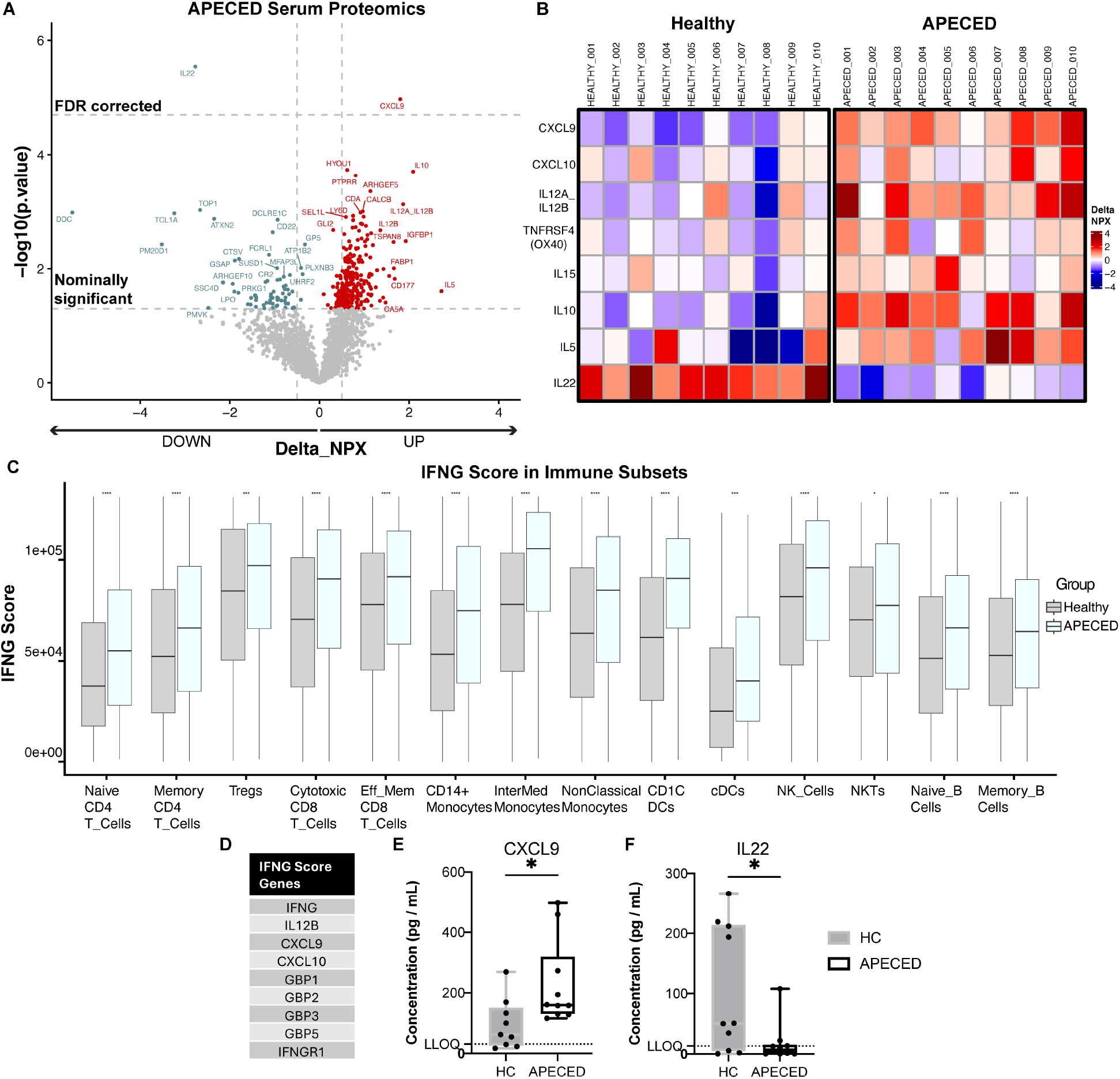
APECED patients have an IFNG skewed inflammatory signature. (A) Serum proteomics in APECED compared to Healthy controls. Average relative protein abundance measured by normalized NPX values (log2 scale) using the Olink platform on X axis. –log10(P-values) from 2 sided welch tests plotted on Y axis with nominal and FDR corrected significance thresholds noted with dotted lines. (B) Healthy and APECED patient serum protein levels for a selection of differentially expressed proteins highlighting a IFNG related signature and cytokine changes. (C-D) (C) Immune cell subsets in Healthy and APECED analyzed with (D) 9 gene IFNG score. Two tailed Wilcoxon rank-sum test with FDR correction. * p < 0.05. (E, F) CXCL9 and IL-22 serum levels validated with secondary assays (Quantikine ELISA and Luminex, respectively). Two tailed Wilcoxon rank-sum test. * p < 0.05.

**Supplemental Figure 6 S6.**
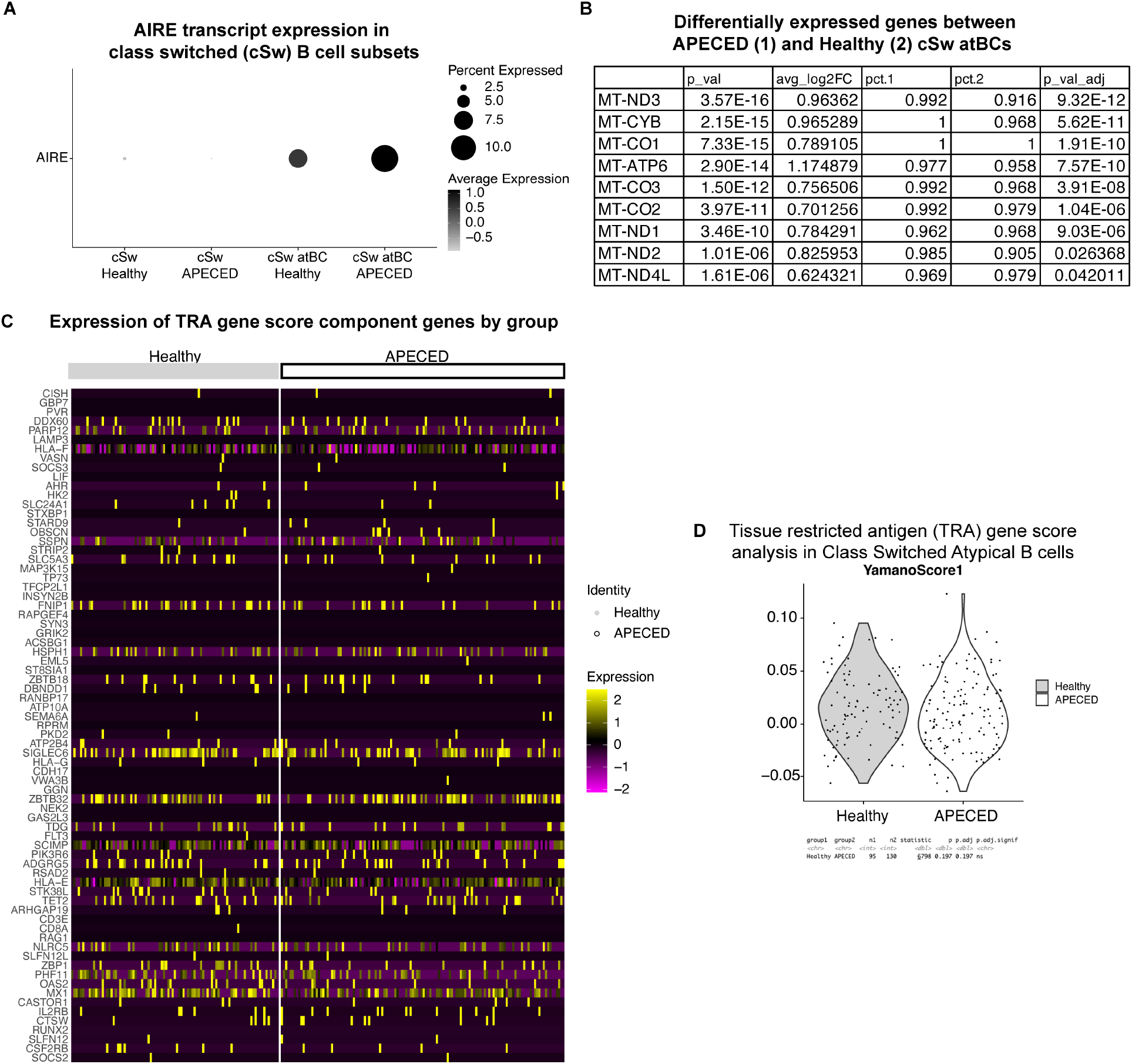
Functional AIRE expression does not significantly impact the transcriptomic state of class switched atypical B cells detected by scRNAseq. (A) AIRE expression in B cell subsets showing expression in cSw atypical B cells. (B) Differential gene expression in cSw atBCs between APECED (group1) and Healthy control (group2). (C) All gene components of the AIRE regulated Tissue restricted antigen (TRA) score from Yamano et al. shown per condition and individual cell. (D) AIRE regulated TRA gene score is not significantly different between Healthy and APECED class switched atypical B cells.

**Supplemental Table 1.**
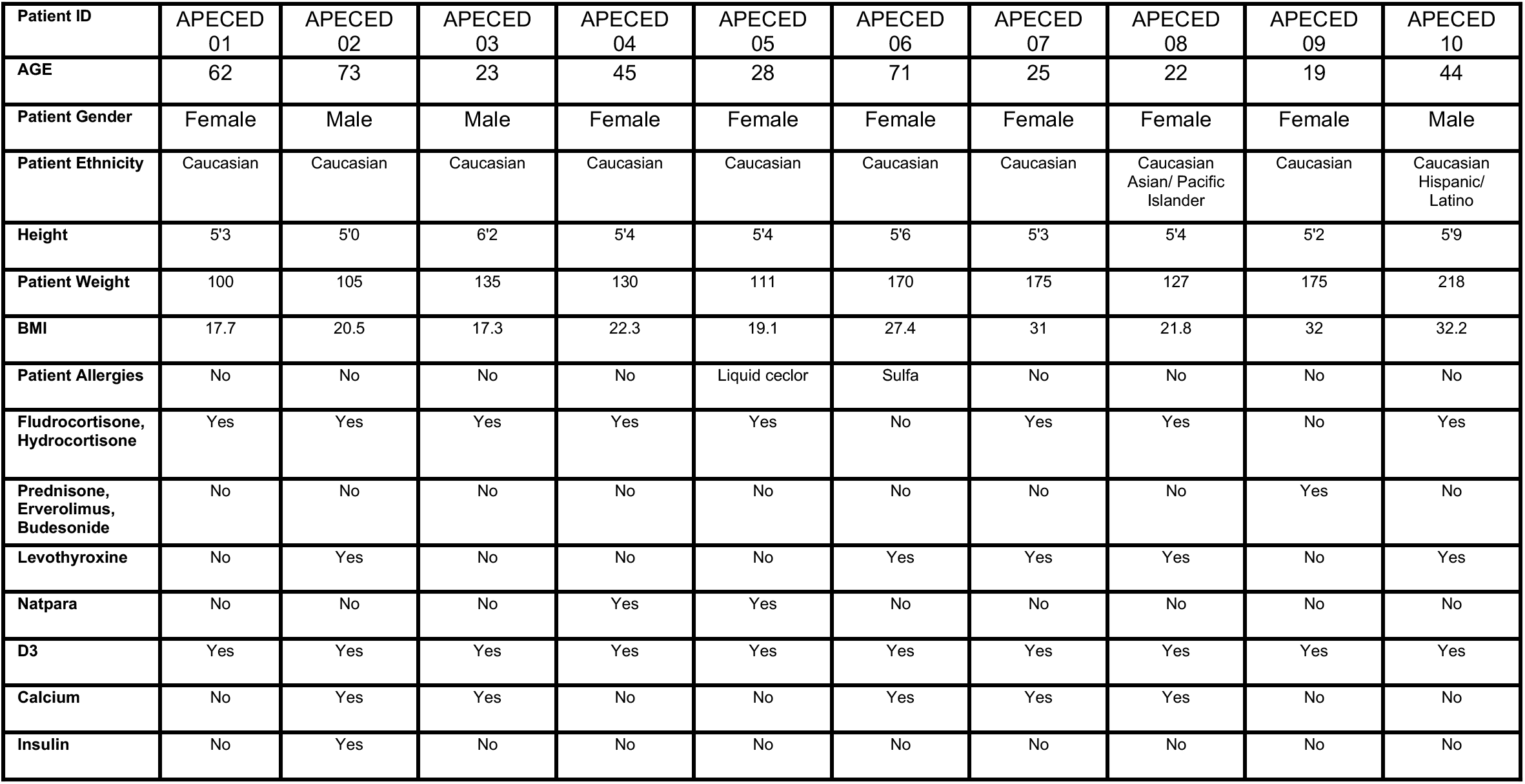
American Autoimmune polyendocrinopathy-Candidiasis-Ectodermal dystrophy (APECED) patient cohort metadata and medications.

**Supplemental Table 2.**
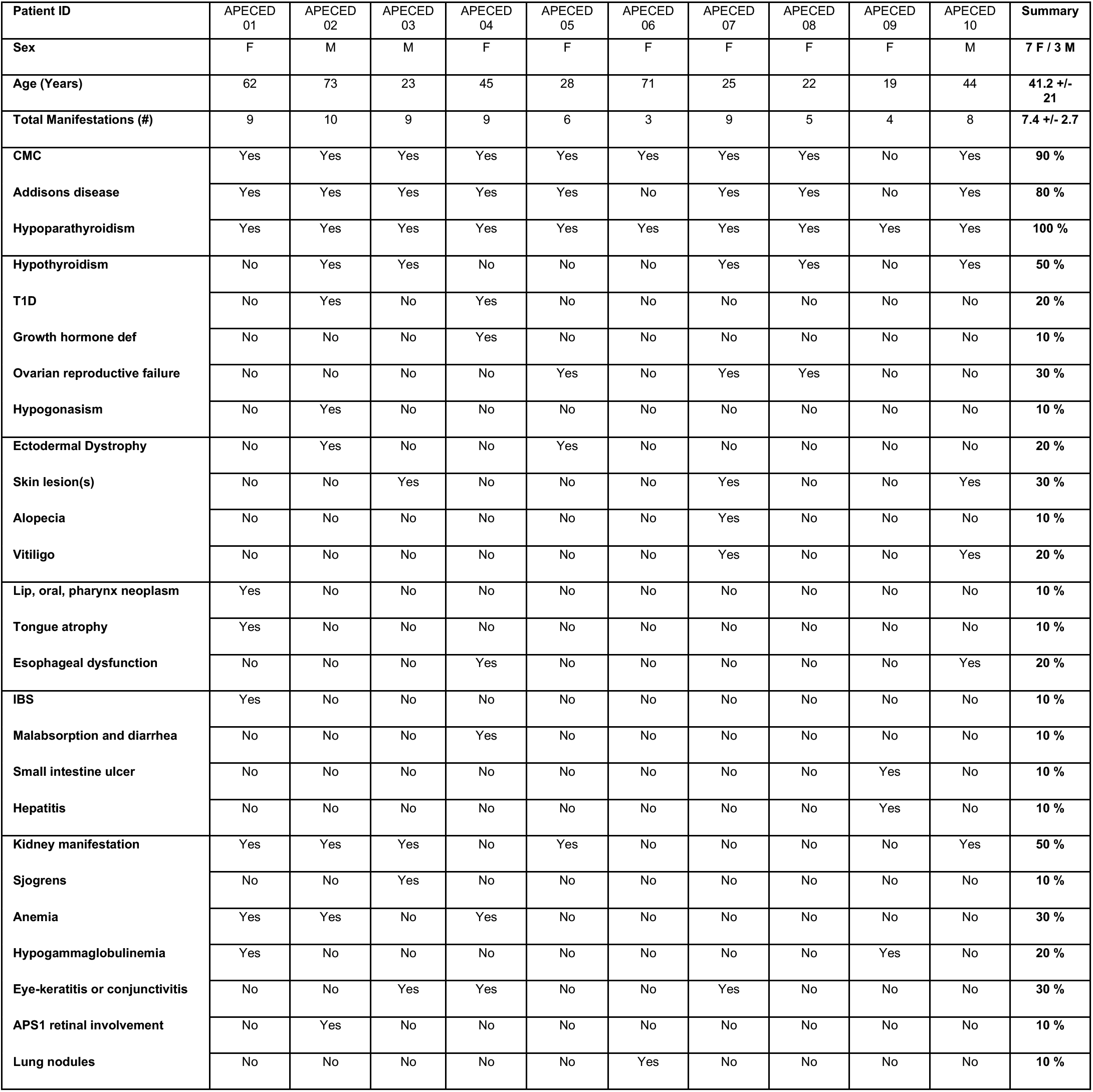
Distribution of clinical manifestations in the 10 Autoimmune polyendocrinopathy-Candidiasis-Ectodermal dystrophy (APECED) patient cohort.

| Patient ID | APECED 01 | APECED 02 | APECED 03 | APECED 04 | APECED 05 | APECED 06 | APECED 07 | APECED 08 | APECED 09 | APECED 10 | Summary |
| --- | --- | --- | --- | --- | --- | --- | --- | --- | --- | --- | --- |
| Sex | F | M | M | F | F | F | F | F | F | M | 7 F / 3 M |
| Age (Years) | 62 | 73 | 23 | 45 | 28 | 71 | 25 | 22 | 19 | 44 | 41.2 +/- 21 |
| Total Manifestations (#) | 9 | 10 | 9 | 9 | 6 | 3 | 9 | 5 | 4 | 8 | 7.4 +/- 2.7 |
| CMC | Yes | Yes | Yes | Yes | Yes | Yes | Yes | Yes | No | Yes | 90 % |
| Addisons disease | Yes | Yes | Yes | Yes | Yes | No | Yes | Yes | No | Yes | 80 % |
| Hypoparathyroidism | Yes | Yes | Yes | Yes | Yes | Yes | Yes | Yes | Yes | Yes | 100 % |
| Hypothyroidism | No | Yes | Yes | No | No | No | Yes | Yes | No | Yes | 50 % |
| T1D | No | Yes | No | Yes | No | No | No | No | No | No | 20 % |
| Growth hormone def | No | No | No | Yes | No | No | No | No | No | No | 10 % |
| Ovarian reproductive failure | No | No | No | No | Yes | No | Yes | Yes | No | No | 30 % |
| Hypogonadism | No | Yes | No | No | No | No | No | No | No | No | 10 % |
| Ectodermal Dystrophy | No | Yes | No | No | Yes | No | No | No | No | No | 20 % |
| Skin lesion(s) | No | No | Yes | No | No | No | Yes | No | No | Yes | 30 % |
| Alopecia | No | No | No | No | No | No | Yes | No | No | No | 10 % |
| Vitiligo | No | No | No | No | No | No | Yes | No | No | Yes | 20 % |
| Lip, oral, pharynx neoplasm | Yes | No | No | No | No | No | No | No | No | No | 10 % |
| Tongue atrophy | Yes | No | No | No | No | No | No | No | No | No | 10 % |
| Esophageal dysfunction | No | No | No | Yes | No | No | No | No | No | Yes | 20 % |
| IBS | Yes | No | No | No | No | No | No | No | No | No | 10 % |
| Malabsorption and diarrhea | No | No | No | Yes | No | No | No | No | No | No | 10 % |
| Small intestine ulcer | No | No | No | No | No | No | No | No | Yes | No | 10 % |
| Hepatitis | No | No | No | No | No | No | No | No | Yes | No | 10 % |
| Kidney manifestation | Yes | Yes | Yes | No | Yes | No | No | No | No | Yes | 50 % |
| Sjogrens | No | No | Yes | No | No | No | No | No | No | No | 10 % |
| Anemia | Yes | Yes | No | Yes | No | No | No | No | No | No | 30 % |
| Hypogammaglobulinemia | Yes | No | No | No | No | No | No | No | Yes | No | 20 % |
| Eye-keratitis or conjunctivitis | No | No | Yes | Yes | No | No | Yes | No | No | No | 30 % |
| APS1 retinal involvement | No | Yes | No | No | No | No | No | No | No | No | 10 % |
| Lung nodules | No | No | No | No | No | Yes | No | No | No | No | 10 % |

